# Ribosome inhibition by *C9ORF72*-ALS/FTD-associated poly-PR and poly-GR proteins revealed by cryo-EM

**DOI:** 10.1101/2020.08.30.274597

**Authors:** Anna B. Loveland, Egor Svidritskiy, Denis Susorov, Soojin Lee, Alexander Park, Gabriel Demo, Fen-Biao Gao, Andrei A. Korostelev

## Abstract

Toxic dipeptide repeat (DPR) proteins are produced from expanded G_4_C_2_ hexanucleotide repeats in the *C9ORF72* gene, which cause amyotrophic lateral sclerosis (ALS) and frontotemporal dementia (FTD). Two DPR proteins, poly-PR and poly-GR, repress cellular translation but the molecular mechanism remains unknown. Here we show that poly-PR and poly-GR of ≥ 20 repeats inhibit the ribosome’s peptidyl-transferase activity at nanomolar concentrations, comparable to specific translation inhibitors. High-resolution cryo-EM structures reveal that poly-PR and poly-GR block the polypeptide tunnel of the ribosome, extending into the peptidyl-transferase center. Consistent with these findings, the macrolide erythromycin, which binds in the tunnel, competes with the DPR proteins and restores peptidyl-transferase activity. Our results demonstrate that strong and specific binding of poly-PR and poly-GR in the ribosomal tunnel blocks translation, revealing the structural basis of their toxicity in *C9ORF72*-ALS/FTD.

## Introduction

Expanded G_4_C_2_ repeats in *C9ORF72* are the most common genetic cause of amyotrophic lateral sclerosis (ALS) and frontotemporal dementia (FTD) (DeJesus-Hernandez et al., 2011; Renton et al., 2011). Unaffected individuals typically carry 5-10 repeats (DeJesus-Hernandez et al., 2011), whereas *C9ORF72*-ALS/FTD patients have from 20-24 (Chen et al., 2016; Gomez-Tortosa et al., 2013; Millecamps et al., 2012) to hundreds or thousands of repeats (Gitler and Tsuiji, 2016; Van Mossevelde et al., 2017). Both sense and antisense repeat RNAs are translated to produce several dipeptide repeat (DPR) proteins in patient neurons (Ash et al., 2013; Mori et al., 2013; Zu et al., 2013). Two of them—poly-PR and poly-GR—are highly toxic in cellular and animal models (Choi et al., 2019; Kwon et al., 2014; Lopez-Gonzalez et al., 2016; Mizielinska et al., 2014; Wen et al., 2014; Yang et al., 2015; Zhang et al., 2018; Zhang et al., 2019). Expression of poly-PR and poly-GR correlates with neurodegeneration in human patient brains (Saberi et al., 2018; Sakae et al., 2018) and results in neurodegeneration in animal models (Hao et al., 2019; Zhang et al., 2018).

Interactome analyses found preferential binding of poly-PR and poly-GR to the translational machinery (Boeynaems et al., 2017; Hartmann et al., 2018; Kanekura et al., 2016; Lee et al., 2016; Lin et al., 2016; Lopez-Gonzalez et al., 2016; Radwan et al., 2020; Tao et al., 2015; Yin et al., 2017), suggesting that these DPR proteins may compromise ribosome functions in patient neurons. Indeed, poly-PR and poly-GR repress translation in mouse and cellular models (Kanekura et al., 2018; Kanekura et al., 2016; Lee et al., 2016; Moens et al., 2019; Zhang et al., 2018) and colocalize with ribosomes in patient brain tissues (Hartmann et al., 2018). However, the molecular mechanism underlying translation repression by poly-PR and poly-GR remains unknown.

## Results

Poly-PR and poly-GR impair global cellular translation (Hartmann et al., 2018; Kanekura et al., 2018; Kanekura et al., 2016; Lee et al., 2016; Moens et al., 2019; Zhang et al., 2018) and associate with the translational machinery *in trans* (Boeynaems et al., 2017; Kanekura et al., 2016; Lin et al., 2016; Yin et al., 2017). To understand how poly-PR and poly-GR disrupt translation, we first used mammalian cell lysates to translate nanoluciferase or firefly luciferase — in the absence or presence of increasing concentrations of PR_20_ and GR_20_ (*the subscript denotes the number of PR or GR repeats*). Both DPR proteins strongly repress translation (**Fig. 1a-b, Supplementary Fig. 1a-b)** with half-maximal inhibition at 300-400 nM (**Fig. 1c-d, Supplementary Fig. 1a-b**), consistent with previous studies showing inhibition of translation in cellular lysates by poly-PR and poly-GR (Kanekura et al., 2018; Kanekura et al., 2016). By contrast, 10 μM poly-GP (glycine-proline)— another soluble product of the G_4_C_2_ expansion — does not inhibit translation (**Supplementary Fig. 1c**). Translation was also not inhibited by 100 μM L-arginine, indicating that the covalent linkage between arginines are required to affect translation (**Supplementary Fig. 1d).** Moreover, poly-PR and poly-GR do not interfere with luciferase enzyme activity or aggregate luciferase mRNAs at these inhibitory concentrations (**Supplementary Fig. 1e-g, Methods**). Thus, poly-PR and poly-GR specifically inhibit translation in a mammalian cell lysate, as strongly as specific ribosome-binding antibiotics, such as harringtonine and cycloheximide (Lando et al., 1976; Lessard and Pestka, 1972; Tscherne and Pestka, 1975).

**Figure 1.**
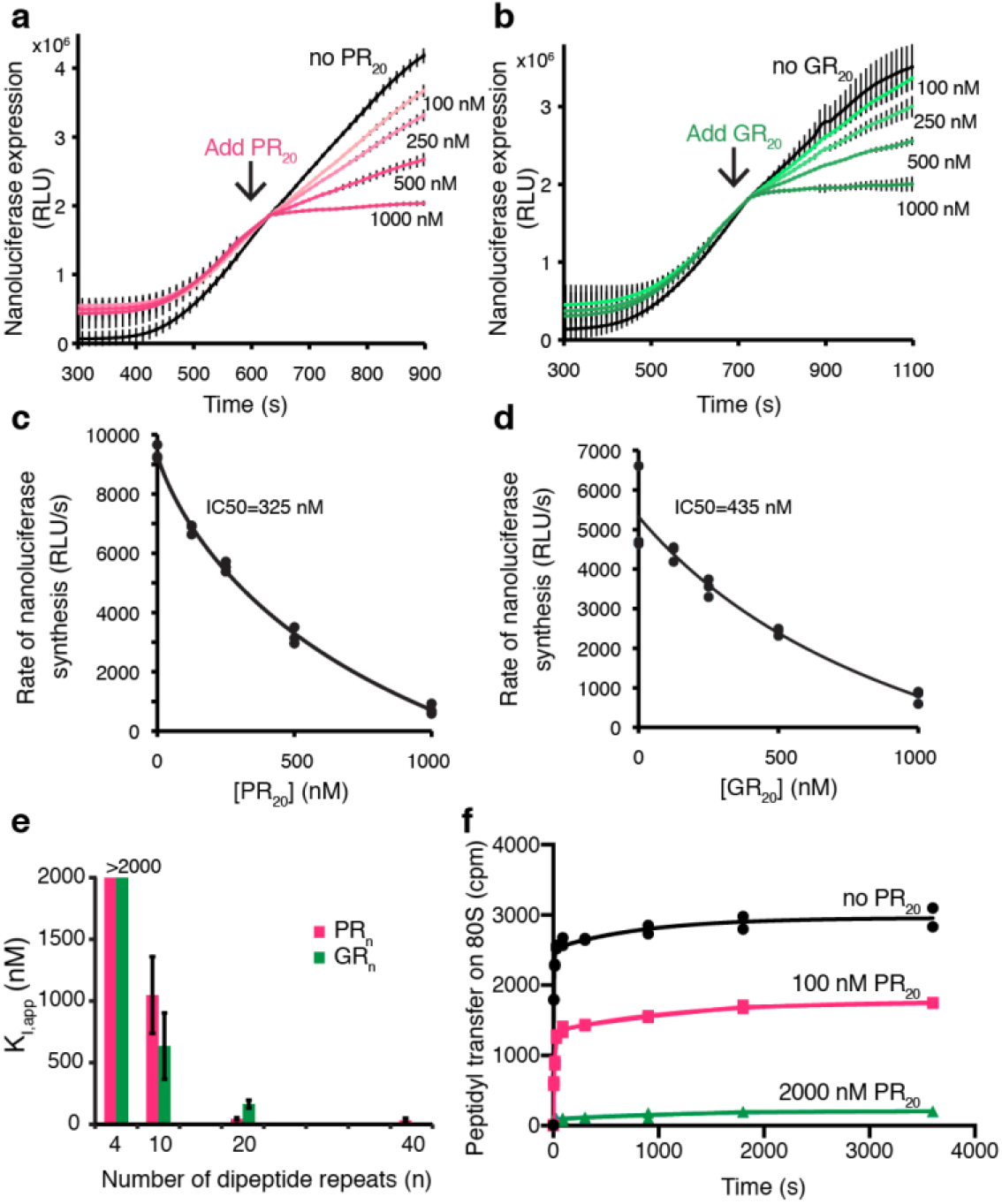
Poly-PR and Poly-GR inhibit translation and peptidyl transfer. (**a-b**) Time courses of the PR_20_ (**a**) and GR_20_ (**b**) inhibition of nanoluciferase mRNA translation in rabbit reticulocyte lysate (RRL; n=3). (**c-d**) Dependence of inhibition of nanoluciferase translation in RRL on the concentrations of PR_20_ and GR_20_ (hyperbola fitting and half-maximal inhibition are shown; n=3). (**e**) Dependence of inhibition of peptide bond formation on the lengths of poly-PR and poly-GR repeats (puromycin assay on *E. coli* ribosomes; apparent inhibition constants are shown on the *y* axis; n=2) (**f**) Concentration dependence of inhibition of peptide bond formation by PR_20_ (puromycin assay on rabbit 80S ribosomes; time progress curves are shown; n=2).

To test if strong translational repression is due to the direct inhibition of the central ribosomal function of peptide bond formation, we measured ribosome-catalyzed peptidyl transfer from peptidyl-tRNA to puromycin (an aminoacyl-tRNA mimic; **Methods**) *in vitro*. Since the arginine-containing DPRs interfere with mitochondrial (Lopez-Gonzalez et al., 2016) and cytoplasmic translation machinery, we measured the inhibition of peptidyl transfer on bacterial 70S ribosomes, a robust system for *in vitro* kinetic studies (Svidritskiy et al., 2013; Wohlgemuth et al., 2006), and mammalian cytoplasmic ribosomes. While L-arginine does not affect the peptidyl transfer reaction (**Supplementary Fig. 1h-i**), PR_20_ and GR_20_ strongly inhibit the reaction with apparent inhibition constants (K_i_) of 44 ± 9 nM and 164 ± 31 nM, respectively (**Fig. 1e**), on 70S ribosomes. The longer PR_40_ confers stronger inhibition (K_i_ ≤30 nM). By contrast, shorter peptides GR_10_ and PR_10_ exhibit a weaker inhibitory effect (K_i_ of 0.6 ± 0.3 μM and 1.0 ± 0.3 μM, respectively). High concentrations of short peptides PR_4_ and GR_4_ (2 μM) fail to inhibit peptidyl transfer (**Supplementary Fig. 1j-k**).

On mammalian ribosomes, PR_20_ starts to inhibit peptidyl transfer at ~100 nM, comparable to that of mammalian ribosome inhibitors (Tscherne and Pestka, 1975) (**Fig. 1f**). GR_20_ is also inhibitory, albeit the inhibition at 2 μM is less than that for PR_20_ (**Supplementary Fig. 1l**). The longer PR_40_ abolishes peptidyl transfer (K_i_ ≤30 nM), whereas the shorter PR_10_ confers no inhibition at 2 μM (**Supplementary Fig. 1m-n**). Thus, the efficiency of inhibition increases with the length of DPR proteins, consistent with a disease-causing length threshold. Our observation of inhibited peptidyl transfer on both bacterial 70S and mammalian 80S ribosomes suggests that longer poly-PR and poly-GR strongly bind a conserved, functionally critical region of the translation machinery.

To understand the structural mechanism of DPR-mediated translation inhibition, we obtained near-atomic resolution cryo-EM structures of PR_20_ and GR_20_ bound to eukaryotic 80S ribosomes (yeast and rabbit) and bacterial 70S ribosomes (**Fig. 2–3**, **Supplementary Fig. 2–3**). The most resolved 2.4-Å structure of the yeast 80S•tRNA•PR_20_ complex reveals poly-PR bound to the polypeptide tunnel, with tRNA excluded from the peptidyl transferase center (PTC) (**Fig**. **2a-c, Supplementary Fig. 2a,c**). The poly-PR chain traverses the tunnel with its N-terminus directed toward the 60S exit. Nine dipeptide repeats of poly-PR are resolved from the PTC through the polypeptide tunnel constriction at the universally conserved 25S rRNA residue A2404 (A3908 in *H. sapiens*; A2062 in *E. coli*) (**Fig. 2b**) toward a second constriction between A883 (A1600 in *H. sapiens*; A751 in *E. coli*) and protein uL4 (**Fig. 2c**), and then toward eL39 and uL22. Arginine and proline residues of PR_20_ stack against 25S rRNA nucleotides, or aromatic or arginine residues of uL4 (**Fig. 2b**, **Supplementary Fig. 2h**), in keeping with strong binding of poly-PR. Several arginine residues are also stabilized by negatively-charged phosphates of 25S rRNA.

**Figure 2.**
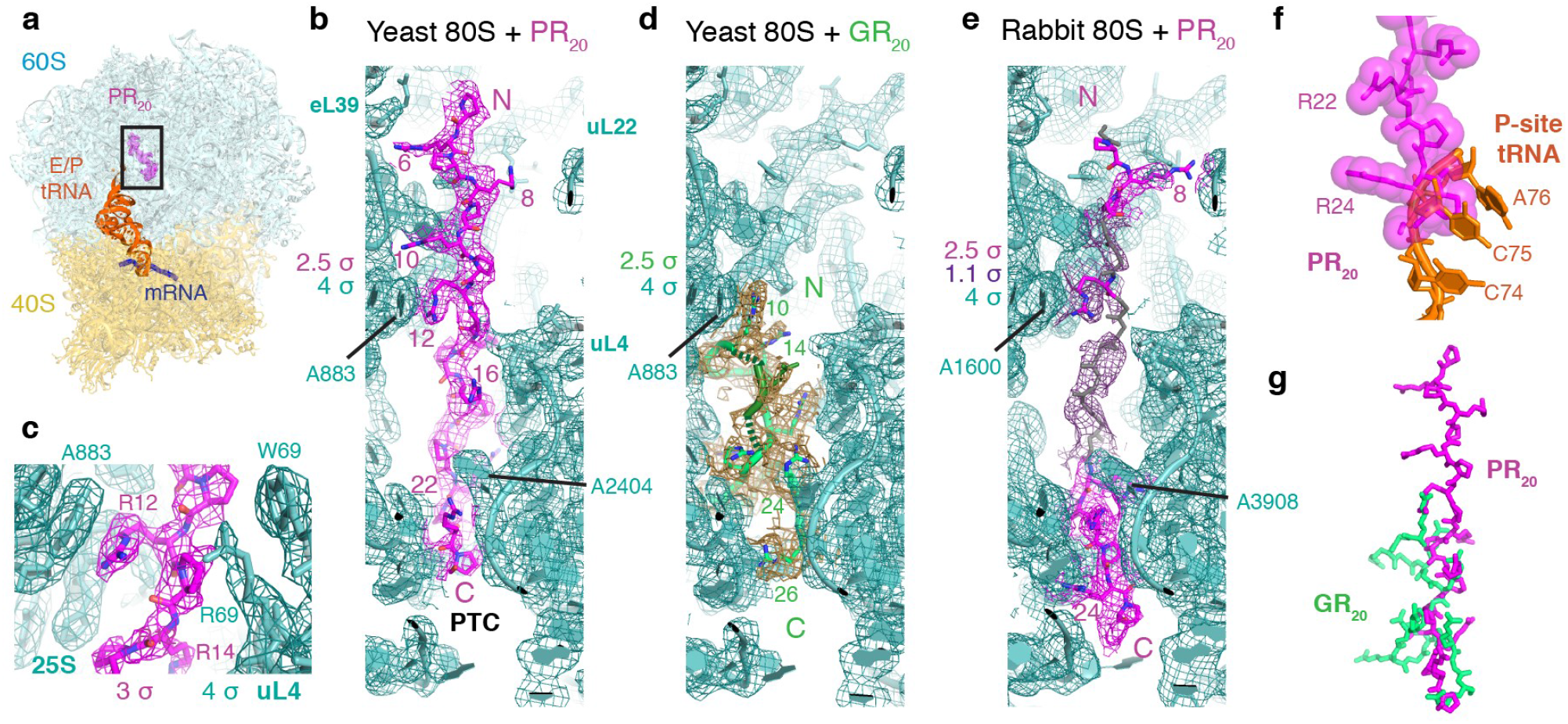
PR_20_ and GR_20_ bind the polypeptide tunnels of eukaryotic 80S ribosomes. (**a**) Overview of the yeast 80S•tRNA•PR_20_ structure. (**b**) Cryo-EM density (mesh) for poly-PR (magenta) in the 80S tunnel (cyan); see also **Supplementary Fig. 2.** Peptidyl transferase center (PTC), ribosomal proteins and nucleotides, and PR repeats are labeled. (**c**) Strong PR density at the constriction at nucleotide A883. (**d**) Cryo-EM density for poly-GR (green) anchored by the tunnel constrictions and adopting alternate conformations (dashed line, dark green) due to glycine flexibility. (**e**) Rabbit 80S•tRNA•PR_20_ map shows strong features (magenta mesh) for PR repeats in PTC and near uL22, connected by lower-resolution density (purple mesh, (see *Methods*)), for which the backbone is modeled (gray). (**f**) Steric clash between poly-PR and superimposed peptidyl-tRNA (orange tRNA 3’ end, from PDB:5LZS, superimposed via 28S rRNA (Shao et al., 2016)). (**g**) Poly-GR makes a more winding path through the tunnel than the relatively straight rod of poly-PR.

**Figure 3.**
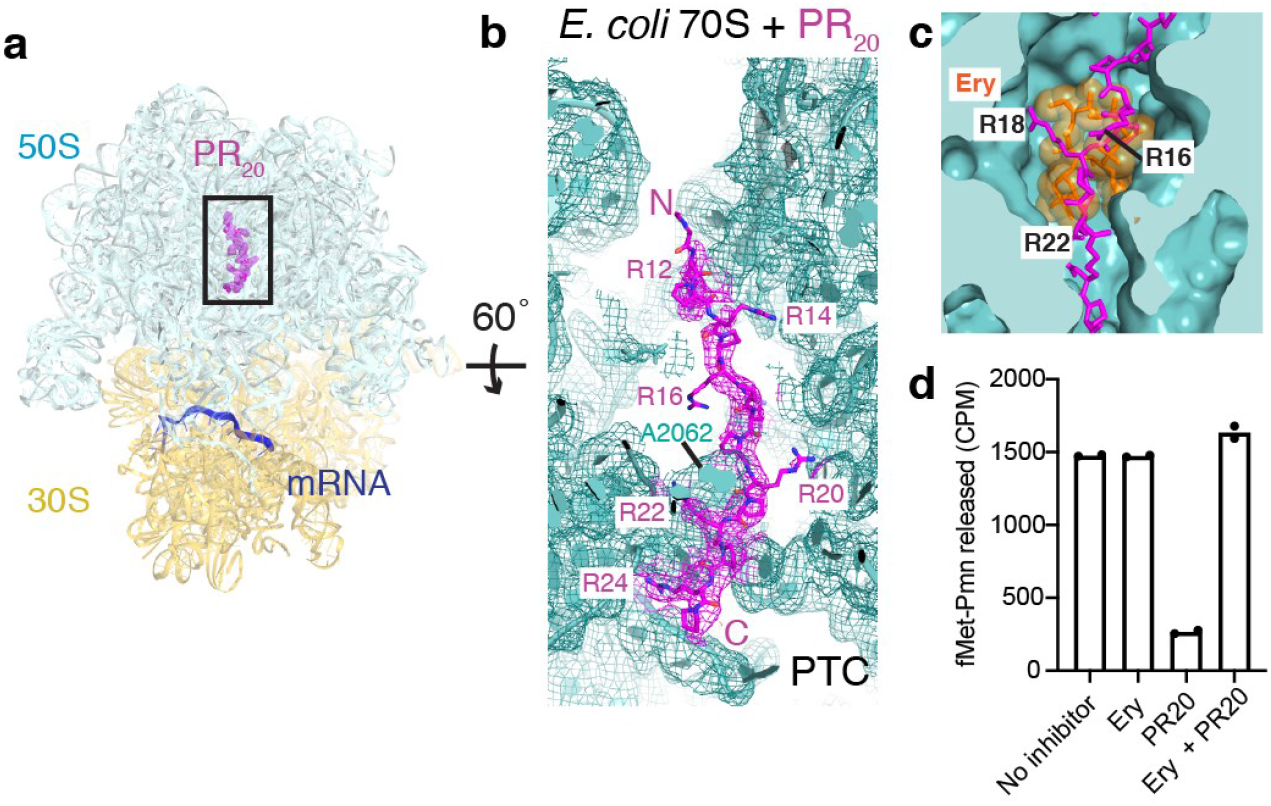
Cryo-EM shows that PR_20_ occupies the polypeptide tunnel of the bacterial 70S ribosome. (**a**) Overview of the structure of *E. coli* 70S•PR_20_. (**b**) PR_20_ binds to the 70S polypeptide exit channel. (**c**) PR_20_ overlaps with the macrolide antibiotic binding site in the polypeptide tunnel. Erythromycin (Ery; orange) from the *E. coli* 70S•Ery crystal structure is shown (PDB:4V7U (Dunkle et al., 2010)). (**d**) Erythromycin (Ery) relieves the inhibition of peptide bond formation by 2 μM PR_20_ indicating an overlapping binding site for Ery and PR_20_.

In the peptidyl transferase center, poly-PR would clash with an amino-acyl or peptidyl moiety on P-site tRNA (**Fig. 2b)**. Lower-resolution features continue away from the PTC toward uL16 and into the intersubunit space typically occupied by A-site tRNA during translation (**Supplementary Fig. 2f**). In the opposite direction—toward the 60S subunit solvent-exposed surface—the widening polypeptide tunnel also shows lower-resolution features corresponding to a continuous poly-PR chain (**Supplementary Fig. 2f**). Together, the ordered and less ordered regions account for ~15 PR repeats, visualizing how the longer poly-PR chains threads the tunnel and inhibits translation by preventing A- and P-tRNA binding in the PTC.

In the 2.7-Å cryo-EM structure of the yeast 80S•tRNA•GR_20_ complex, the most interpretable features of poly-GR are also located between the PTC and the tunnel constrictions (**Fig. 2d, Supplementary Fig. 2b-c**) where poly-GR takes a winding path, reaching deep into crevices of the polypeptide tunnel (**Fig. 2g)**. Similar to the poly-PR complex, lower-resolution features of GR_20_ extend to the tunnel exit on the 60S surface (**Supplementary Fig. 2g**). The features of poly-GR, however, are less resolved than those of poly-PR, reflecting the flexibility of glycine residues.

The structure of the polypeptide tunnel, where poly-PR and poly-GR bind, is nearly fully conserved between yeast and mammals **(Supplementary Fig. 2h**). Indeed, our 3.1-Å resolution cryo-EM structure of the rabbit 80S•tRNA•PR_20_ complex, though less resolved confirms that poly-PR binds similarly in the mammalian and yeast ribosomes, which feature nearly identical polypeptide tunnels (**Fig. 2e, Supplementary Fig. 2h**). Again, poly-PR density is highest in the tunnel constriction near A3908 of 28S rRNA. Here, poly-PR reaches into the PTC and either competes with P-site tRNA (**Fig. 2f, Supplementary Fig. 2i-j,l**) or packs on the ribose of the 3’ terminal nucleotide of deacylated tRNA (**Supplementary Fig. 2k**, see **Methods**). Thus, poly-PR is incompatible with the methionyl moiety of the initiating tRNA (**Supplementary Fig. 2m**) and with longer peptidyl-tRNA.

Maximum-likelihood classification of our cryo-EM datasets revealed that all yeast and mammalian 80S ribosome states contained poly-PR or poly-GR in the polypeptide tunnel, consistent with the high affinity of these DPR proteins in our biochemical experiments (**Supplementary Fig. 2a,b,i**). No other binding sites were identified. Although we cannot exclude non-specific binding to poorly resolved peripheral regions of the ribosome, these distant interactions are unlikely to strongly interfere with peptidyl transfer. Importantly, the classifications also revealed that the DPR proteins bind the polypeptide tunnels of isolated yeast and mammalian 60S large ribosomal subunits (**Supplementary Fig. 2d,e,l**). Thus, poly-PR and poly-GR binding does not depend on the small 40S subunit and may affect the steps before peptidyl transfer, which include 60S subunit maturation and subunit association during translation initiation (**see Discussion**).

Finally, cryo-EM analyses of *E. coli* 70S ribosomes and 50S subunits confirm that PR_20_ is stably held at the constrictions of the conserved polypeptide tunnel to inhibit peptidyl transfer (**Fig. 3a-b, Supplementary Fig. 3, Supplementary Information**). Here, we formed an initiation 70S complex with fMet-tRNA^fMet^ and performed an EF-Tu•GTP-catalyzed elongation reaction with Val-tRNA^Val^ and EF-Tu•GTP, in the presence of PR_20_. Cryo-EM resolved several elongation states of the 70S ribosome at up to 2.9-Å average resolution, as discussed in (**Supplementary Information**). The high abundance of vacant 70S with PR_20_ (**Fig. 3a, Supplementary Fig. 3b-c**) indicates that poly-PR inhibits formation of the initiation 70S•fMet-tRNA^fMet^ complex. Moreover, we observed tRNA-bound initiation and elongation states were sampled almost exclusively by the ribosomes that lacked poly-PR, consistent with the incompatibility of poly-PR with formation of the peptide bond (**Supplementary Fig. 3e-g**). The structures of the PTC and tunnel constrictions of the *E. coli* ribosome are nearly identical to those of mammalian mitochondrial ribosomes (Aibara et al., 2020; Amunts et al., 2015; Brown et al., 2017; Greber et al., 2015; Greber et al., 2014; Koripella et al., 2020; Kummer and Ban, 2020; Kummer et al., 2018) (**Supplementary Fig. 3j**). Indeed, mitochondrial ribosomes are sensitive to antibacterial ribosome inhibitors that bind in this region (de Vries et al., 1973; Ibrahim et al., 1974). Although we have not tested mitochondrial translation, the structural and functional conservation suggests that arginine-containing DPR proteins could bind and inhibit mitochondrial ribosomes, as supported by the association of cellular DPR proteins with mitochondrial ribosome components (Hartmann et al., 2018; Lopez-Gonzalez et al., 2016; Radwan et al., 2020).

We next asked whether the inhibition of peptidyl transfer is directly caused by binding of the DPR proteins in the polypeptide tunnel, rather than by a non-specific mechanism, e.g. binding of the positively-charged DPR proteins to ribosomal, transfer or messenger RNAs. We employed a competition assay, using the macrolide antibiotic erythromycin, an inhibitor of bacterial translation. Erythromycin binds at the constriction of the bacterial polypeptide tunnel near the peptidyl-transferase center (**Fig. 3c**), but does not perturb the puromycin reaction (Cundliffe and McQuillen, 1967). Thus, erythromycin is expected to restore peptidyl-transferase activity if it displaces poly-PR from the tunnel. We find that in the presence of both erythromycin and inhibitory PR_20_, the peptide bond formation for fMet-puromycin is restored (**Fig. 3d**), indicating that inhibition of peptidyl transfer by poly-PR is due to polypeptide tunnel binding.

## Discussion

This work uncovers how toxic arginine-containing DPR proteins inhibit ribosome function. Our structures and biochemical results show that longer poly-PR and poly-GR strongly bind *in trans* to the polypeptide tunnels of all ribosome species we tested, from bacterial to mammalian, as well as of the isolated large ribosomal subunits to repress translation (**Fig. 4**). Poly-PR and poly-GR binding to large subunits and ribosomes strongly interferes with initiator Met-tRNA binding and peptidyl transfer, consistent with translation initiation (Moens et al., 2019) and elongation defects (Kanekura et al., 2018; Kanekura et al., 2016) (**Fig. 4a**). By contrast, several other arginine-rich polypeptides, whose arginine content ranges from 46% to 100%, do not to cause translation inhibition or cell death (Kanekura et al., 2018), emphasizing the specificity of poly-PR and poly-GR action. Poly-PR and poly-GR inhibition therefore results from strong and specific binding to the polypeptide tunnel via electrostatic and packing interactions, which are most pronounced for poly-PR (**Figs. 2, 3**). Our biochemical and structural findings are consistent with the higher cellular toxicity of poly-PR than poly-GR (Hartmann et al., 2018; Kanekura et al., 2018; Kwon et al., 2014; Lee et al., 2016; Wen et al., 2014). Our data are also consistent with the disease-causing length threshold of DPR proteins (Chen et al., 2016; Gomez-Tortosa et al., 2013; Millecamps et al., 2012), revealing that the longer DPR proteins traverse the tunnel constrictions into the PTC, protruding into the tRNA binding sites (**Supplementary Fig. 2f)**.

**Figure 4.**
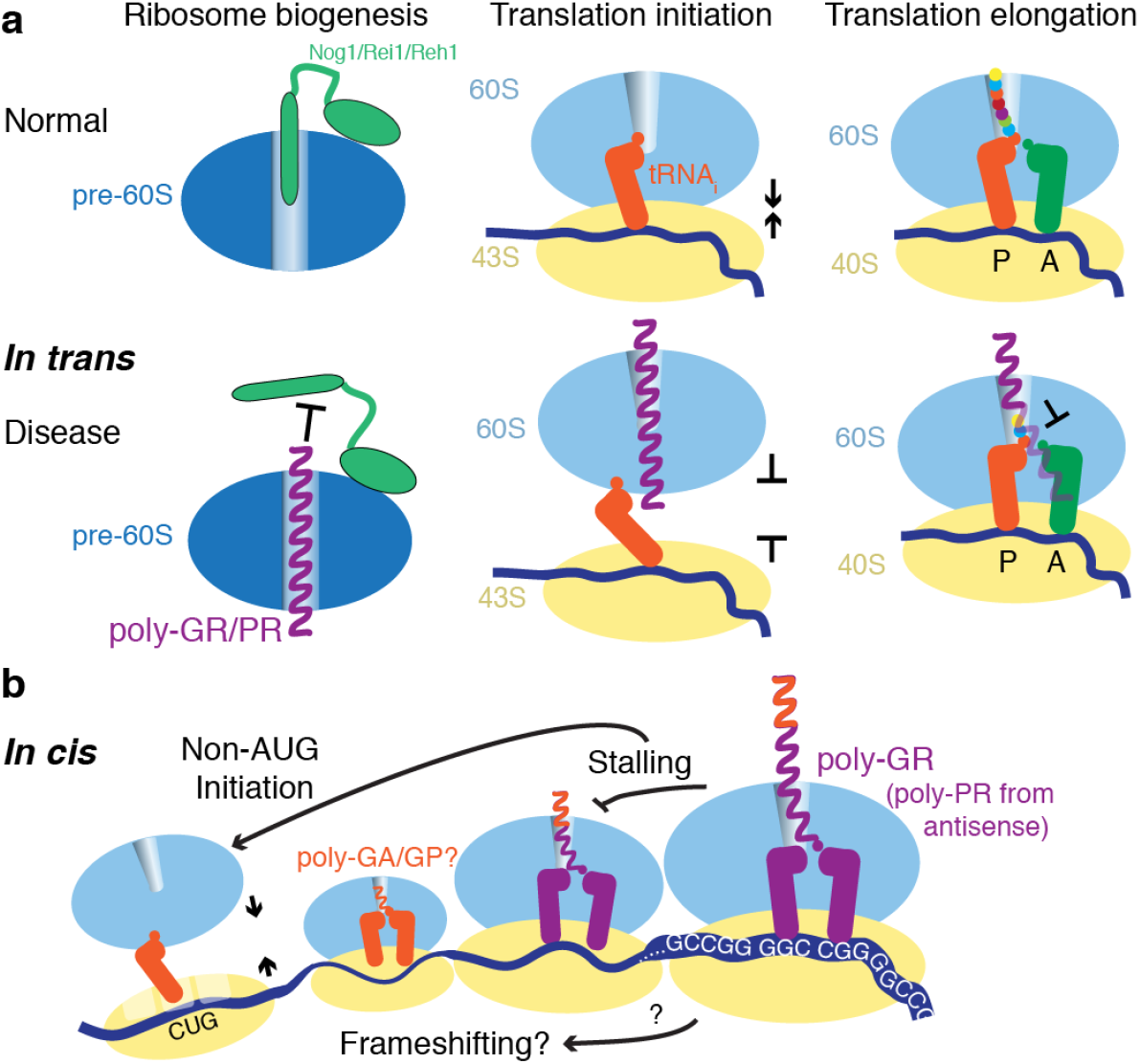
Proposed mechanisms of translation repression caused by poly-PR and poly-GR binding to the ribosomal polypeptide tunnel. (**a**) *In trans* binding of poly-PR and poly-GR (magenta) to 60S (blue) polypeptide tunnel is consistent with perturbation of 60S biogenesis (Hartmann et al., 2018; Kwon et al., 2014; Wen et al., 2014) and translation initiation (Moens et al., 2019), while binding to 80S inhibits translation elongation (this work and (Kanekura et al., 2016)). Similarly, binding to large mitochondrial subunits and mitochondrial ribosomes may perturb mitochondrial function (Choi et al., 2019; Lopez-Gonzalez et al., 2016). Biogenesis factors Nog1/Rei1/Reh1 are green, 40S subunit yellow, tRNAs in the P and A sites are orange and green. (**b**) Translation of G_4_C_2_ repeats may cause stalling of poly-PR and poly-GR in the ribosome, resulting in translation inhibition *in cis*. If transient, the stalling may enhance otherwise inefficient non-AUG initiation and/or frameshifting (Kearse et al., 2019) of the G_4_C_2_ repeat sequence (Tabet et al., 2018), leading to amplified translation of the DPR proteins (tRNA/peptide is orange for non-arginine DPR proteins and magenta for arginine-containing DPR proteins).

Our analyses were initially inspired by our finding that poly-GR preferentially binds to mitochondrial and cytoplasmic ribosomal proteins (Lopez-Gonzalez et al., 2016). Remarkably, the mode of poly-PR and poly-GR action resembles proline-rich antimicrobial peptides (PrAMPs), produced by eukaryotes as an antimicrobial strategy (Graf and Wilson, 2019). The PrAMPs share a conserved PRP repeat sometimes expanded to (XR)_4_, which binds at the polypeptide tunnel similarly to poly-PR and poly-GR in our work (**Supplementary Fig. 4a-d**). PrAMPs also inhibit translation and compete with macrolide antibiotics for binding (Florin et al., 2017a; Gagnon et al., 2016b; Mardirossian et al., 2018; Seefeldt et al., 2016; Seefeldt et al., 2015), echoing our findings with erythromycin (**Fig. 3e**). The ability of mammalian cells to express PrAMPs (Mardirossian et al., 2018) demonstrates that the inhibitory repeat sequences with (XR)n repeats can be produced by cytoplasmic ribosomes and then interfere with translation *in trans*.

Indeed, cellular co-localization of the arginine-containing DPR proteins with nucleoli (Hartmann et al., 2018; Kwon et al., 2014; Wen et al., 2014) and mitochondrial components (Hartmann et al., 2018; Lopez-Gonzalez et al., 2016; Radwan et al., 2020), and the extracellular localization of DPRs (Wen et al., 2014) demonstrate that *C9ORF72*-translated poly-PR and poly-GR are liberated from ribosomes into the cytoplasm and organelles, and impairing global translation. The inefficient translation required to make DPR proteins (Radwan et al., 2020) may delay their accumulation to toxic levels, in keeping with the late onset of ALS and FTD. Although concentrations of soluble DPR proteins in individual patient neurons are not known and difficult to measure, we speculate that accumulation of poly-GR and/or poly-PR in some neurons may be facilitated by stress conditions and/or processes associated with aging. The accumulating DPR proteins likely bind the ribosomal subunits and ribosomes, and gradually dysregulate translation, leading to neuron degradation.

Our findings also suggest how poly-PR and poly-GR could interfere with other cellular processes involving the ribosome, which are beyond the scope of this study but supported by published research. First, consistent with cellular co-localization of the arginine-containing DPR proteins with nucleoli in cell lines and patient tissues (Hartmann et al., 2018; Kwon et al., 2014; Lee et al., 2016; Tao et al., 2015; Wen et al., 2014), impaired ribosomal rRNA biogenesis in cells expressing DPR proteins (Kwon et al., 2014; Tao et al., 2015), and the identification of ribosomal biogenesis factors as DPR interactors (Kanekura et al., 2016; Lee et al., 2016; Radwan et al., 2020; Tao et al., 2015), DPR binding to the maturing pre-60S particles might disrupt ribosome biogenesis. Indeed, the DPR protein binding site directly overlaps with ribosome biogenesis factors Nog1, Rei1, and Reh1, which enter the ribosomal polypeptide tunnel to test the integrity of the maturing 60S subunit (**Fig. 4a, Supplementary Fig. 4e**) (Greber et al., 2016; Ma et al., 2017; Wu et al., 2016). Next, the structural and functional conservation suggests that arginine-containing DPRs could bind and inhibit mitochondrial ribosomes, consistent with the association of cellular DPR proteins with mitochondrial ribosome components (Hartmann et al., 2018; Lopez-Gonzalez et al., 2016; Radwan et al., 2020) and mitochondrial dysfunction associated with *C9ORF72* repeat expansions (Choi et al., 2019; Lopez-Gonzalez et al., 2016). Finally, poly-GR and/or poly-PR could affect translation of extended G_4_C_2_- or G_2_C_4_-repeat RNA. Here, nascent poly-GR and poly-PR could bind tightly to the polypeptide tunnel and stall ribosomes *in cis* (**Fig. 4b**). Indeed, when we finalized this work, a study reported that poly-PR and poly-GR but not poly-GA or poly-AP stall translation *in cis*, requiring more than 10 repeats for efficient stalling (Radwan et al., 2020). Co-translational stalling may result in accumulation of DPR-stalled polysomes (Hartmann et al., 2018; Yamada et al., 2019), consistent with DPR- and ribosome-containing aggregates in patient cells (Hartmann et al., 2018; Mori et al., 2013), and trigger cellular stress (Tao et al., 2015). Increased cellular stress may enhance DPR protein synthesis (Cheng et al., 2018; Green et al., 2017), eliciting a positive feedback loop between DPR protein accumulation and cellular stress. Indeed, translational stalling was recently shown to enhance translation from non-AUG codons (Kearse et al., 2019). Since translation of G_4_C_2_ mRNA repeats depends on non-AUG initiation and possibly frameshifting (Tabet et al., 2018), translation of arginine-containing DPR proteins may increase cellular production of these and/or other DPR proteins (Yamada et al., 2019), such as poly-GA and poly-GP.

Our work does not exclude other — ribosome-independent—mechanisms that may contribute to *C9ORF72*-ALS/FTD, including: defective nuclear transport (Shi et al., 2017; Vanneste et al., 2019), defective splicing (Kwon et al., 2014; Yin et al., 2017), aggregation of non-arginine DPRs, such as poly-GA (Guo et al., 2018), repeat-RNA toxicity, or loss of functional *C9ORF72* protein and more (as reviewed in refs (Balendra and Isaacs, 2018; Gitler and Tsuiji, 2016)). Yet, concrete structural mechanisms of macromolecular inhibition for these possible pathways remain to be determined. Our findings of strong atomic interactions between the arginine-containing DPR proteins and the ribosome, and impairment of the central ribosome function, suggest a direct structural mechanism for cellular toxicity in *C9ORF72*-ALS/FTD and possibly in spinocerebellar ataxia type 36 (SMA36) where accumulation of poly-PR has recently been observed (McEachin et al., 2020).

## Acknowledgements

We thank Chen Xu, KangKang Song, and Kyounghwan Lee for help with screening cryo-EM grids and for assistance with data collection at the UMass Medical School cryo-EM facility; Christine Carbone for assistance with ribosome purification; Darryl Conte Jr. and members of the Korostelev lab for helpful comments on the manuscript. This study was supported by the Dan and Diane Riccio Fund for Neuroscience (to AAK and FBG), and by NIH Grants R01NS101986 (to FBG), and R01GM107465 and R35GM127094 (to AAK).

## Data availability

The models generated and analyzed during the current study will be available from the RCSB Protein Data Bank.

## Methods

GR_10_, GR_20_, GR_40_, PR_4_, PR_10_, PR_20_, PR_40_ peptides were synthesized and purified by CS Bio Co (Menlo Park, CA). Mass-spectrometry and HPLC were used to confirm the length and purity of peptides. Solid peptides were dissolved in water yielding stock concentrations between 1 and 10 mM, which were further diluted to specific concentrations for biochemical and cryo-EM experiments.

### Luciferase assays in Rabbit Reticulocyte Lysates

Rabbit reticulocyte lysates (RRL, Promega) were used to investigate the inhibition of translation by DPR proteins. We independently monitored translation of two mRNAs encoding non-homologous proteins of different lengths: nanoluciferase and firefly luciferase. Aliquots of 10 μl of the reaction mixture, containing 30 mM Hepes-KOH, 50 mM KoAc, 1 mM Mg(OAc)_2_, 0.2 mM GTP and ATP, 2mM DTT, 0.02 mM of each amino acid (Promega), 33% RRL (Promega) and 1% nanoluciferase substrate (Promega), were incubated in 384-well plate (Nunc, white) at 30°C for 5 min inside of microplate reader (Tecan Infinite m1000 pro). Then a model mRNA, coding for nanoluciferase (England et al., 2016) (Promega) was added to the aliquots to the final concentration of 30 nM. Translation reactions were incubated at 30°C for 20 min, luminescence of the samples was continuously measured by the microplate reader to record the time course of protein synthesis. After approximately 10 min from the reaction start the measurement was paused and GR_20_, PR_20_ or cycloheximide were added to the final concentrations of 0-1 μM or 2.5 μM, respectively, then the measurement continued. Time progress curves, obtained in the experiments, were processed in MS Excel, with triplicate measurements normalized to the RLU signal at the point of DPR protein or cycloheximide addition. To obtain inhibition efficiency, kinetic curves were approximated by linear equations over the interval of 100 seconds after DPR protein or water (control) addition. The slopes of the equations were plotted versus DPR protein concentrations, and the plots were fitted by hyperbolas in Gnuplot to obtain the IC50 values. Firefly luciferase translation was performed similarly to that for nanoluciferase, except that the efficiency of translation of 100 ng/μl firefly luciferase mRNA (Promega) in RRL was measured (in relative luminescence units, RLU) after 1 hour of reaction incubation at 30^°^C, in the absence or presence of PR_20_, GR_20_ and GP20 (control) (**Supplementary Fig. 1a-c**).

The following additional controls indicate that DPR-induced changes in fluorescence (**Fig. 1a-c**) were caused by translation inhibition. First, translation was not inhibited by 100 μM L-arginine, indicating that covalent linkages between arginines are required (**Supplementary Fig. 1d**). Second, poly-PR and poly-GR do not disrupt luciferase or nanoluciferase activity, when added after the luciferases have been translated in the absence of poly-PR and poly-GR (**Supplementary Fig. 1e-f**). Finally, they do not aggregate mRNAs at the inhibitory concentrations as described below (**Supplementary Fig. 1g**).

### Measurement of DPR-induced RNA aggregation

To test whether arginine-containing DPR proteins cause mRNA aggregation at the DPR concentrations sufficient to inhibit protein synthesis (0.2 and 1 uM), we measured turbidity (optical density at 600 nm; OD600; similarly to (Kanekura et al., 2016)) of the solutions containing 100 ng/μl total HEK293 RNA. We incubated RNA with PR_20_, GR_20_, GP_20_ (control) or water for 20 minutes at room temperature. Addition of PR_20_ and GR_20_ at concentrations that inhibit RRL translation and peptidyl transfer (from 0.2 to 1 μM) did not increase optical density relative to RNA alone, nor did water or GP_20_. 10 μM PR_20_ and GR_20_ did not induce visible opalescence but resulted in increased OD600 (**Supplementary Fig. 1g**). Optical density was measured using NanoDrop One (Thermo Scientific, Waltham, MA, USA).

### Kinetic peptidyl transfer assays

The effects of GR_n_ or PR_n_ on peptide bond formation in *E. coli* ribosome were tested in an *in vitro* assay that reacts the 70S ribosome loaded with [^35^S]-fMet-tRNA^fMet^ in the P site with puromycin, an A-site tRNA mimic, and measures the release of the resulting dipeptide containing formyl-methionine covalently bonded to puromycin, as described (Svidritskiy and Korostelev, 2018). *E. coli* tRNA^fMet^ (Chemical Block) was aminoacylated with [^35^S]-N-formyl-L-methionine (Perkin Elmer) as described (Lancaster and Noller, 2005). 70S ribosomes were prepared from *E. coli* (MRE600) as described (Svidritskiy and Korostelev, 2018). 70S ribosomes were stored in the ribosome-storage buffer (20 mM Tris-HCl, pH 7.0; 100 mM NH_4_Cl; 20 mM MgCl_2_) at −80°C. A model mRNA fragment, containing the Shine-Dalgarno sequence and a spacer to position the AUG codon in the P site and the UUC codon in the A site (GGC AAG GAG GUA AAA <u>AUG UUC</u> AAAAAA), was synthesized by IDT. 70S•mRNA•fMet-tRNA^fMet^ complexes with GR_n_ (where n = 4, 10 or 20) or PR_n_ (where n = 4, 10, 20 or 40) were formed as follows. 330 nM 70S ribosomes were incubated with 11 μM mRNA in buffer containing 20 mM Tris-acetate (pH 7.0), 100 mM NH_4_OAc, 12 mM Mg(OAc)_2_, for two minutes at 37°C. 730 nM [^35^S]-fMet-tRNA^fMet^ was added and incubated for five minutes at 37°C. The solution was diluted 9.6-fold with the same buffer, followed by addition of a GR_n_ or PR_n_ repeat peptide (22x relative to its final concentration). The complex was incubated for five minutes at 37°C prior to addition of puromycin dissolved in the same buffer (see below).

To test the effects of PR_20_ on the mammalian peptidyl transferase, 80S ribosomes were purified from rabbit reticulocyte lysate (Green Hectares) and dissociated into 40S and 60S subunits as described in the section entitled “Cryo-EM complex of mammalian 80S ribosome with PR_20_.” A model leaderless mRNA fragment placing the AUG codon in the P site and the UUC codon in the A site (CCAC <u>AUG UUC</u> CCCCCCCCCCCCCCCCCC), was synthesized by IDT. The 80S•mRNA•fMet-tRNA^fMet^ complex with PR_n_ (where n=10, 20 or 40) or GR_20_ was assembled as follows. 1 μM 60S subunit was mixed with the DPR (starting with the initial DPR concentration of 33x relative to its final concentration in the reaction) in buffer containing 20 mM Tris-acetate (pH 7.0), 100 mM KOAc, 10 mM Mg(OAc)_2_, and the mixture was incubated for five minutes at 30°C. In a separate microcentrifuge tube, 600 nM 40S was mixed with 16.5 uM mRNA in the same buffer and incubated for two minutes at 30°C. 1095 nM [^35^S]-fMet-tRNA^fMet^ and the 60S solution were added (0.5 volumes of the 40S solution), resulting in the following concentrations: 330 nM 60S, 400 nM 40S, 730 nM tRNA, 11 uM mRNA and 11x of final concentration of a DPR in the buffer(20 mM Tris-acetate (pH 7.0), 100 mM KOAc, 10 mM Mg(OAc)_2_). The solution was incubated for five minutes at 30°C. The solution was diluted 10 times with the same buffer, then puromycin (prepared in the same buffer) was added and time progress curves were recorded.

The kinetics of puromycin reaction on the 70S or 80S ribosomes were recorded and analyzed essentially as described (Svidritskiy et al., 2013). An aliquot (4.5 μl) of the complex prior to addition of puromycin was quenched in 30 μl of saturated MgSO_4_ in 0.1 M NaOAc to represent the zero-time point. 5 μl of 500 μM puromycin (in the same buffer as that used for the ribosomal complex) was added to 45 μl of the complex to initiate the reaction, yielding estimated final concentrations: 30 nM for the ribosome, 1 uM mRNA, 66 nM [^35^S]-fMet-tRNA^fMet^, 50 μM puromycin, and varying concentrations of PR_n_ and GR_n_ peptides, from 30 nM to 2 μM. After 6, 15, 30, 90 seconds, and 5, 15, 30, 60 and (for slower reactions) 180 minutes, 5-μl aliquots were quenched in 30 μl of saturated MgSO_4_ in 0.1 M NaOAc. All quenched samples were extracted with 700 μl ethyl-acetate. 600 μl of the extract were mixed with 3.5 ml of Econo-Safe scintillation cocktail (RPI), and the amount of released [^35^S]-labeled dipeptide was measured for each time point using a scintillation counter (Beckman Coulter, Inc.).

All time progress curves were recorded in duplicates, using independently prepared 70S or 80S complexes. Reaction rates were calculated using single-exponential or double-exponential (80S complexes) regressions for different concentrations of PR_n_ and GR_n_ using GraphPad Prism 8. Hyperbola fitting in Gnuplot was used to derive apparent K_i_ values.

#### Competition assay using erythromycin

was performed as described above for the puromycin kinetics, except that the 70S ribosomes were pre-incubated with 22 μM erythromycin for 5 minutes prior to adding PR_20_. The stock solution of erythromycin contained 2 mM erythromycin in ethanol, so the control reaction (70S with no erythromycin) and the reactions with erythromycin were prepared to contain the same amount of ethanol (1%). The final concentrations after addition of puromycin were: 30 nM 70S, 66 nM [^35^S]-fMet-tRNA^fMet^, 1 μM mRNA, 2 μM PR_20_ (no PR_20_ in the control reaction), 20 μM erythromycin (or no erythromycin in a control reaction), 50 μM puromycin (in 20 mM Tris-acetate (pH 7.0), 100 mM NH_4_OAc, 12 mM Mg(OAc)_2_ and 1% ethanol). Samples were quenched at 300 s when the uninhibited reaction had reached a plateau.

### Cryo-EM complexes of yeast 80S ribosomes with PR_20_ or GR_20_

*Saccharomyces cerevisiae* 80S ribosomes complexes with PR_20_ or GR_20_, were assembled *in vitro* as follows. *S. cerevisiae* 40S and 60S ribosomal subunits were purified from strain W303 and stored in 1x Reassociation Buffer 50 mM Tris-HCl pH 7.5, 20 mM MgCl_2_, 100 mM KCl, 2 mM DTT, as previously described (Abeyrathne et al. 2016). To form the 80S complex, 12 pmol of 40S ribosomal subunits were incubated in 1x Reassociation Buffer with 240 pmol of a custom RNA oligonucleotide (Integrated DNA Technologies, Inc.) encoding the Kozak sequence, the start codon AUG, and an open reading (CCAC-AUG-UUC-CCC-CCC-CCC-CCC-CCC-CCC) and 60 pmol of tRNA^fMet^ (ChemBlock) for 5 minutes at room temperature. Subsequently, 14.3 pmol of 60S ribosomal subunits were added and incubated for a further 5 minutes at room temperature. PR_20_ or GR_20_ were diluted to 30 μM in Buffer B: 50 mM Tris-HCl pH 7.5, 10 mM MgCl_2_, 100 mM KCl and mixed 1:1 with the 80S assembly reaction for a final reaction containing: 300 nM 40S, 360 nM 60S, 6 μM mRNA, 1.5 μM tRNA^fMet^, and 15 μM GR_20_ or 15 μM PR_20_. Complexes were incubated for a further 5-10 minutes at room temperature prior to plunging cryo-EM grids.

Quantifoil R2/1-4C grids coated with a 2-nm thin layer of carbon were purchased from EMSDiasum. The grids were glow discharged with 20 mA current with negative polarity for 60 seconds in a PELCO easiGlow glow discharge unit. A Vitrobot Mark IV (ThermoFisher Scientific) was pre-equilibrated to room temperature and 95% humidity. 2-3 μl of the 80S assembly reaction was applied to the grid, incubated for 10-20 seconds, blotted for 5 seconds, and then plunged into liquid-nitrogen-cooled liquid ethane.

### Cryo-EM complex of mammalian 80S ribosome with PR_20_

*Oryctolagus cuniculus* 80S complex with PR_20_ was prepared using ribosomal subunits purified from rabbit reticulocyte lysate (RRL), based on the procedures described in references (Lomakin and Steitz, 2013; Pisarev et al., 2007). 60 ml of RRL (Green Hectares) was thawed and mixed 1:1 in 2x RRL dissolving buffer (10 mM HEPES pH 7.1, 30 mM KCl, 22 mM Mg(OAc)_2_, 2 mM EDTA, 4 mM DTT, 0.6 mg/ml heparin, and Complete Protease Inhibitor (Roche)). The solution was split and layered over four sucrose cushions of 20 mM Bis-Tris pH 5.9, 300 mM KCl, 200 mM NH_4_Cl, 10 mM Mg(OAc)_2_, 30% (w/v) sucrose and 5 mM DTT and centrifuged in Ti45 rotor for 16 hours at 43,000 rpm. The crude ribosome pellets were gently resuspended in re-dissolving buffer (20 mM Tris-HCl pH 7.5, 50 mM KCl, 4 mM Mg(OAc)_2_, 1 mM EDTA, 2 mM DTT, 1 mg/ml heparin, and Complete Protease Inhibitor (Roche)). To disrupt polysomes, 1 mM puromycin was added to ribosome solution and incubated 20 minutes at 37°C, 20 minutes at room temperature, and 20 minutes on ice. To separate subunits, the KCl concentration in the ribosome solution was gradually adjusted from 50 mM to 500 mM followed by a 30-minute incubation at 4°C. The ribosomal subunits were separated using a 10-35% sucrose gradient (20 mM Tris-HCl pH 7.5, 500 mM KCl, 4 mM Mg(OAc)_2_, 2 mM DTT). 12 tubes were centrifuged for 16 hours at 22,000 rpm in SW28 rotors, and the fractions for the 40S and 60S peaks were collected using a Gradient Master (Biocomp Instruments). The 40S and 60S subunits were concentrated and exchanged into Storage Buffer (20 mM Tris-HCl pH 7.5, 100 mM KCl, 2.5 mM Mg(OAc)_2_, 2mM DTT, 0.25 M sucrose) using centrifugal filters (100 kDa MWCO; Amicon).

The complex of rabbit 80S ribosomes with PR_20_ for cryo-EM was prepared similarly to that for puromycin release assays. 1 μM rabbit 60S subunits were pre-incubated for 5 minutes at 30°C with 10 μM PR_20_ in buffer (20 mM Tris-acetate (pH 7.0), 100 mM KOAc, 10 mM Mg(OAc)_2_). Meanwhile, 40S subunits were preincubated with mRNA for 5 minutes at 30°C in the same buffer. tRNA^fMet^ (Chemical Block) was added to 40S such that final concentrations were 1.2 μM 40S, 40 μM mRNA, and 2.4 μM tRNA^fMet^ and then the 40S and 60S reactions were mixed 1:1 and incubated for 5 minutes at 30°C. The complex was then diluted 60% with the same buffer and PR_20_ and placed on ice until cryo-plunging. The final complex contained: 200 nM 60S, 240 nM 40S, 8 μM mRNA, 480 nM tRNA^fMet^, and 5 μM PR_20_.

400M copper grids coated with lacey carbon and a 2-nm thin layer of carbon (Ted Pella Inc.) were glow discharged with 20 mA current with negative polarity for 60 s in a PELCO easiGlow glow discharge unit. A Vitrobot Mark IV (ThermoFisher Scientific) was pre-equilibrated to 4°C and 95% relative humidity. 2.5 μl of the 80S assembly reaction was applied to grid, incubated for 10 seconds, blotted for 4 seconds, and then plunged into liquid-nitrogen-cooled liquid ethane.

### Cryo-EM complex of bacterial 70S ribosome with PR_20_

30S and 50S ribosomal subunits were prepared from MRE600 *E. coli* as described (Svidritskiy and Korostelev, 2018) and stored in Buffer A (20 mM Tris-HCl, pH 7, 10.5 mM MgCl_2_, 100 mM NH_4_Cl, 0.5 mM EDTA, 6 mM β-mercaptoethanol) at −80°C. *E. coli* EF-Tu, fMet-tRNA^fMet^ and Val-tRNA^Val^ (tRNA^Val^ from Chemical Block) were prepared as described (Loveland et al., 2017). mRNA containing the Shine-Dalgarno sequence and a linker to place the AUG codon in the P site and the Val codon (cognate complex) in the A site was synthesized by Integrated DNA Technologies Inc (GGC AAG GAG GUA AAA <u>AUG</u> <u>GUA</u> AGU UCU AAA AAA AAA AAA).

The 70S complexes were prepared as follows. Heat-activated (42°C, 5 minutes) 30S ribosomal subunits (1 μM) were mixed with 50S ribosomal subunits (1 μM) and with mRNA (4 μM) (all final concentrations) in Reaction Buffer (20 mM HEPES•KOH, pH 7.5, 20 mM MgCl_2_, 120 mM NH_4_Cl, 2 mM spermidine, 0.05 mM spermine, 2 mM β-mercaptoethanol) for 10 minutes at 37°C. Equimolar fMet-tRNA^fMet^ was added to the ribosomal subunits and incubated for 3 minutes at 37°C. Subsequently, 15 μM PR_20_ was added and incubated another 5 minutes at 37°C, then the reaction was held on ice. Concurrently, the ternary complex of Val-tRNA^Val^•EF-Tu•GTP was prepared as follows. 6 μM EF-Tu was pre-incubated with 1 mM GTP (Roche) in Reaction Buffer for 5 minutes at 37°C and then was supplemented with 4 μM Val-tRNA^Val^ (all final concentrations). After an additional minute at 37°C, the ternary complex reaction was also kept on ice to prepare for plunging. The 70S reaction and ternary complex was mixed in 1x Reaction Buffer and incubated at 37°C for 10 minutes for the elongation step to complete. The final reaction had the following concentrations: 330 nM 50S; 330 nM 30S; 1.3 μM mRNA; 330 nM fMet-tRNA^fMet^; 500 nM EF-Tu; 83 μM GTP, 330 nM Val-tRNA^Val^ with or without 5 μM PR_20_.

Quantifoil R2/1 holey-carbon grids coated with a thin layer of carbon (EMSDiasum) were glow discharged with 20 mA current with negative polarity for 60 seconds in a PELCO easiGlow glow discharge unit. A Vitrobot Mark IV (ThermoFisher Scientific) was pre-equilibrated to 5°C and 95% relative humidity. 2.5 μl of the ribosomal complex prepared with 5 μM PR_20_ were applied to chilled grids, and then blotted for 4 seconds prior to plunging into liquid-nitrogen-cooled liquid ethane.

### Electron Microscopy

Data for the *S. cerevisae* 80S•PR_20_ or 80S•GR_20_ were collected on a Titan Krios electron microscope (ThermoFisher Scientific) operating at 300 kV and equipped with a Gatan Image Filter (Gatan Inc.) and a K2 Summit direct electron (Gatan Inc.) targeting 0.5 to 2.0-μm underfocus. For 80S•PR_20_, a dataset of 203,089 particles from 3033 movies was collected automatically using SerialEM (Mastronarde, 2005) using beam tilt to collected 5 movies per hole at 4 holes between stage movements (Svidritskiy et al., 2019). The movies had a total of 30 frames with 1 e^−^/Å^2^ per frame for a total dose of 30 e^−^/Å^2^ on the sample. The 80S•GR_20_ dataset had 467,615 particles from 3451 movies collected using beam tilt to collected 6 shots per hole. 2071 movies were used for data analysis after exclusion of suboptimal movies (due to poor contrast, ice or collection outside of desired area). The movies had a total of 32 frames with 0.9 e^−^/Å^2^ per frame for a total dose of 30 e^−^ /Å^2^ on the sample. The super-resolution pixel size was 0.5294 Å for both datasets.

Data for two datasets- (1) rabbit 80S•PR_20_ (2) *E. coli* 70S•PR_20_ —were collected on a Talos electron microscope (ThermoFisher Scientific) operating at 200 KV and equipped with a K3 direct electron detector (Gatan Inc.) targeting 0.6 to 1.8-μm underfocus. Data collection was automated using SerialEM (Mastronarde, 2005) using beam tilt to collect multiple movies (e.g. 4 movies per hole at 4 holes) at each stage position (Svidritskiy et al., 2019). The *E. coli* dataset had a total of 20 frames per movie, with 1.1 e^−^/Å^2^ per frame for a total dose of 36 e^−^/Å^2^ on the sample, comprising 1024 movies yielding 172,278 particles for the 70S•PR_20_ complex. The rabbit 80S dataset had 3031 movies, at 19 frames per movie at 1.5 e^−^/Å^2^ per frame for a total dose of 30 e^−^/Å^2^ on the sample, yielding 246,885 particles. Movies were aligned on the fly during data collection using IMOD (Kremer et al., 1996) to decompress frames, apply the gain reference, and to correct for image drift and particle damage yielding image sums with pixel size of 0.87 Å (K3).

### Cryo-EM data classification and maps

#### Yeast 80S complexes

Early steps of 3D map generation from CTF determination, reference-free particle picking, and stack creation were carried out in cisTEM, while particle alignment and refinement was carried out in Frealign 9.11 (Lyumkis et al., 2013b) and cisTEM (Grant et al., 2018). To speed up processing, 2×-, 4×- and 6×-binned image stacks were prepared using resample.exe, which is part of the Frealign distribution (Lyumkis et al., 2013a). The initial model for particle alignment of 80S maps was EMD-5976 (Svidritskiy et al., 2014), which was down-sampled to match 6× binned image stack using EMAN2 (Tang et al., 2007). Three rounds of mode 3 search alignment to 20 Å were run using the 6× binned stack. Next, 25-30 rounds of mode 1 refinement were run with the 4×, 2× and eventually unbinned stack until the resolutions stopped improving (2.96 Å for PR_20_ and 3.06 Å for GR_20_), similarly to the “auto-refine” procedure in cisTEM. Next, one round of beam shift refinement and per-particle CTF refinement improvement the final maps to 2.41 Å (PR_20_) and 2.73 Å (GR_20_), which were used for model building. 3D maximum likelihood classification into 6 classes was used to separate the 60S maps shown in **Supplementary Fig. 2d-e**.

To separate the ribosomes with and without DPR proteins in the tunnel, we used maximum-likelihood classification into up to 6 classes, applying a 30 A focus mask around the center of the polypeptide tunnel of the 2× stack at the resolution of 6 Å. This strategy revealed features for poly-PR or poly-GR in all classes, but the density differed suggesting that heterogeneous conformations of the chains are possible in the tunnel.

#### Rabbit 80S complex

CTF determination, micrograph screening to remove off-target shots, reference-free particle picking, and stack creation were carried out in cisTEM. Particle alignment and refinement was carried out in Frealign 9.11 and cisTEM. To speed up processing, 4×- and 8×-binned image stacks were prepared using resample.exe. The initial model for particle alignment of 80S maps was EMD-4729, which was downsampled to match the 8× binned stack and low-pass filtered to 30 Å, using EMAN2. Two rounds of mode 3 search alignment to 20 Å were performed using the 8× binned stack. Next, 7 rounds of mode 1 refinement were run with the 4× and eventually the unbinned stack as we gradually added resolution shells (limit of 6 Å) and map resolution reached 3.50 Å. Next, one round of beam shift refinement improved the resolution to 3.11 Å and one round of per-particle CTF refinement (to 6 Å) improved the resolution to 3.02 Å. 3D maximum likelihood classification into 12 classes was then used to resolve different ribosome states including the free 60S state (3.5 A) (**Supplementary Fig. 2i**).

Maximum-likelihood classification revealed 80S ribosomes with a single P/E tRNA (rotated 80S states) and with two tRNAs bound in the P and E sites (non-rotated 80S states), yielding different poly-PR density features in the PTC (**Supplementary Fig. 2i-l**). The 3.1-Å map used for poly-PR modeling and structure refinement was reconstructed from three classes corresponding to the 80S ribosome with a hybrid-state P/E tRNA (>50% occupancy and score >0). This conformation is overall similar to the rotated conformation of yeast 80S ribosomes with a single P/E tRNA (**Fig. 2a**). Here, poly-PR traverses the PTC and is incompatible with the binding of aminoacylated (**Supplementary Fig. 2m**) or deacylated P-tRNA (**Fig. 2f**, P-site tRNA is shown from the structure of rabbit 80S elongation complex with amino-acyl-tRNA, eEF1A, and didemnin PDB:5LZS (Shao et al., 2016) after superposition of the 28S rRNA). By contrast, non-rotated 80S states had an additional tRNA in the E site, forcing the deacylated P-tRNA into the 60S P site (**Supplementary Fig. 2i**). In these maps, poly-PR extends toward the CCA end of the deacylated P-site tRNA (**Supplementary Fig. 2k**) and is incompatible with an aminoacyl-tRNA (as in the initiator Met-tRNA) or peptidyl-tRNA (**Supplementary Fig. 2m**). The particles comprising the 4 classes of the 80S ribosome with P/P and E/E tRNA were merged into a single map (**Supplementary Fig. 2i, k**).

In the isolated 60S subunit density (3.5 Å resolution), poly-PR density is similar to that in the rotated 80S ribosome (**Supplementary Fig. 2l**).

In **Fig. 2e**, the map is shown for the rotated 80S state, with ribosomal residues at 4σ (cyan) and PR residues at 2.5σ (magenta) within the PTC and near uL22. Continuous lower-resolution density between these regions is shown using a low-pass-filtered map (to 4 Å, B-factor softened by applying a B-factor of 50) at 1.1σ (dark purple).

#### 70S complex

Early steps of 3D map generation from CTF determination, reference-free particle picking, and stack creation were carried out in cisTEM, while particle alignment and refinement was carried out in Frealign 9.11 and cisTEM. To speed up processing, 2×-, and 8×-binned image stacks were prepared using resample.exe. The initial model for particle alignment of 70S maps was the 11.5 Å map EMD-1003 (Gabashvili et al., 2000), which was down-sampled to match the 8× binned image stack using EMAN2. 3 rounds of mode 3 search aligned to 20 Å were run using the 8× binned stack. Next, multiple rounds of mode 1 refinement were run with the 8×, 2× stacks, and eventually the unbinned stack, as we gradually added resolution shells (limit of 6 Å) and resolution was 2.98 Å (PR_20_). Next, beam shift refinement was used to improve the overall resolution to 2.86 Å, also resulting in improved map features for both rRNA and proteins.

3D maximum likelihood classification was used to separate particles into classes of different compositions (**Supplementary Fig. 3b**). Classes were added until classification revealed the hybrid 70S class with the P/E tRNA and A/P tRNA (the product of the dipeptide reaction). This processing required 12 classes for 60 rounds for the 70S•PR_20_ complex. PR_20_ classes were reconstructed with the original refined parameters (to 6 Å maximum resolution) and overall beam shift parameters to yield the final map for model building (obtained by merging four 70S classes without tRNA) or structural analyses (**Supplementary Fig. 3c-h**). Comparison of polyPR and erythromycin binding sites in **Fig. 3c** was prepared by structural superimposition of the 23S rRNA of the 70S•tRNA•PR_20_ structure to that of *E. coli* ribosome bound with erythromycin (Dunkle et al., 2010).

### Model building and refinement

#### Yeast 80S complexes

The 3.0-Å crystal structure of the *Saccharomyces cerevisiae* 80S ribosome (PDB: 4V88 (Ben-Shem et al., 2011)) was used as a starting model for structure refinement. Because of the heterogeneity in the density for the 40S subunit conformation in our highest-resolution map, we only modeled and refined the 60S subunit and the tRNA bound to the L1 stalk. The model for the P/E-site tRNA^fMet^ and the L1 stalk was derived from the 6.2-Å cryo-EM structure of the *Saccharomyces cerevisiae* 80S ribosome bound with 1 tRNA (PDB: 3J77 (Svidritskiy et al., 2014)). PR_20_ was modeled in Coot(Emsley and Cowtan, 2004) in either C-out or N-out conformations starting from the PrAMPs Bac7 (PDB:5HAU (Gagnon et al., 2016b)) or Api137 (PDB:5O2R (Florin et al., 2017a)) and their fits into well-resolved density near residue A883. Local real-space refinement in Phenix (Afonine et al., 2018) with and without Ramachandran restraints was used to determine the best directionality (N-out) for the PR chain in the polypeptide exit channel. GR_20_ was similarly modeled in Coot and refined.

#### Rabbit 80S complex

The 3.0-Å cryo-EM structure of XBP1u-paused *Oryctolagus cuniculus* ribosome-nascent chain complex (PDB: 6R5Q (Shanmuganathan et al., 2019)) was used as a starting model for structure refinement. The 60S subunit was refined into the 3.1 Å cryo-EM map with PR_20_, which was remodeled in Coot, starting with the best fitting N-out conformation from the yeast 80S-PR_20_ complex.

#### 70S complex

The 3.2-Å cryo-EM structure of 70S•tRNA•EF-Tu•GDPCP (PDB: 5UYM (Loveland et al., 2017)) was used as a starting model for structure refinements with tRNAs and EF-Tu removed. The ribosome was fitted into maps using Chimera (Pettersen et al., 2004) with independent fitting for the 50S, L1 stalk, L11 stalk, 30S body, shoulder, and head. Local adjustments to nucleotides in the polypeptide exit channel and decoding center were made using Pymol (DeLano, 2002). PR_20_ was remodeled in Coot, starting from the N-out conformations from the yeast 80S-PR_20_ complex.

#### Model refinement

The models were refined using real-space simulated-annealing refinement using RSRef (Chapman, 1995; Korostelev et al., 2002) against corresponding maps. Refinement parameters, such as the relative weighting of stereochemical restraints and experimental energy term, were optimized to produce the optimal structure stereochemistry, real-space correlation coefficient and R-factor, which report on the fit of the model to the map (Zhou et al., 1998). The structures were next refined using phenix.real_space_refine (Adams et al., 2011) to optimize protein geometry, followed by a round of refinement in RSRef to optimize RNA geometry, applying harmonic restraints to preserve protein geometry (Chapman, 1995; Korostelev et al., 2002). Phenix was used to refine B-factors of the models against their respective maps (Adams et al., 2011). The resulting structural models have good stereochemical parameters, characterized by low deviation from ideal bond lengths and angles and agree closely with the corresponding maps as indicated by high correlation coefficients and low real-space R factors (**Table S1**). Figures were prepared in Pymol and Chimera.

---

**Supplementary Fig. 1.**
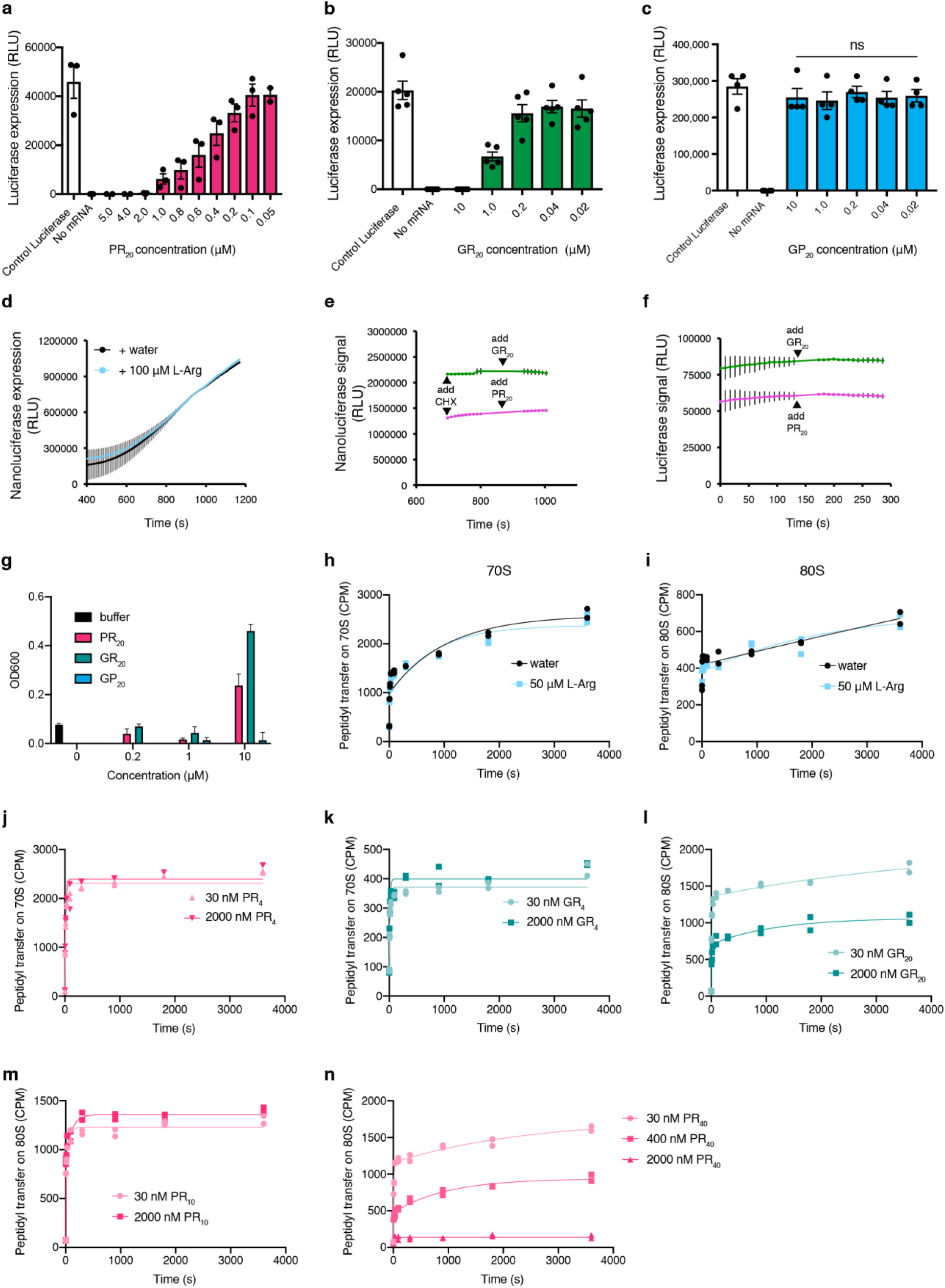
Control experiments for inhibition of translation by DPRs. (**a-c**) PR_20_ and GR_20_ but not GP_20_ inhibit translation of firefly luciferase mRNA in RRL. Relative luminescence units (RLU) after 1 hour of translation in the absence (positive luciferase control and negative “no mRNA” control) or presence of DPRs are shown; n=3, 5 and 4, respectively. (**d**) 100 μM L-Arg does not inhibit the translation of nanoluciferase in RRL. (**e-f**) PR_20_ and GR_20_ do not affect enzyme activity of nanoluciferase (**e**) and firefly luciferase (**f**). Nanoluciferase mRNA was translated in the absence of DPRs, translation was stopped by cycloheximide (CHX), then 1 μM PR_20_ or GR_20_ were added and luminescence was monitored. 40 nM firefly luciferase enzyme (Promega) was supplemented into RRL in the absence of mRNA and amino acids, and after 120 seconds 1 μM PR_20_ or GR_20_ were added. n=3 (**g**) Concentrations of PR_20_ and GR_20_ that inhibit RRL translation and peptidyl transfer (from 0.2 to 1 μM) do not cause HEK293 RNA aggregation nor does water or GP_20_, measured as 600 nm light absorbance. 10 μM PR_20_ and GR_20_ did not induce visible opalescence but result in increased OD600, consistent with some RNA aggregation (n=3, error bars are SD). (**h-i**) Time progress curves of the puromycin reaction showing that 50 μM L-Arg (blue squares) does not inhibit peptidyl transfer on *E. coli* 70S ribosomes (**h**) and rabbit 80S ribosomes (**i**). (**j-k**) Time progress curves of the puromycin reaction on *E. coli* 70S ribosomes showing that PR_4_ and GR_4_ do not inhibit peptidyl transfer at high (2 μM) concentration. (**l**) Time progress curves of the puromycin reaction on rabbit 80S ribosomes showing that GR_20_ inhibits peptidyl transfer at 2 μM. (**m**) Time progress curves of the puromycin reaction on rabbit 80S ribosomes showing PR_10_ does not inhibit peptidyl transfer at 2 μM. (**n**) Time progress curves of the puromycin reaction on rabbit 80S ribosomes showing PR40 strongly inhibits peptidyl transfer (n=2 in panels **h-n**).

**Supplementary Fig. 2:**
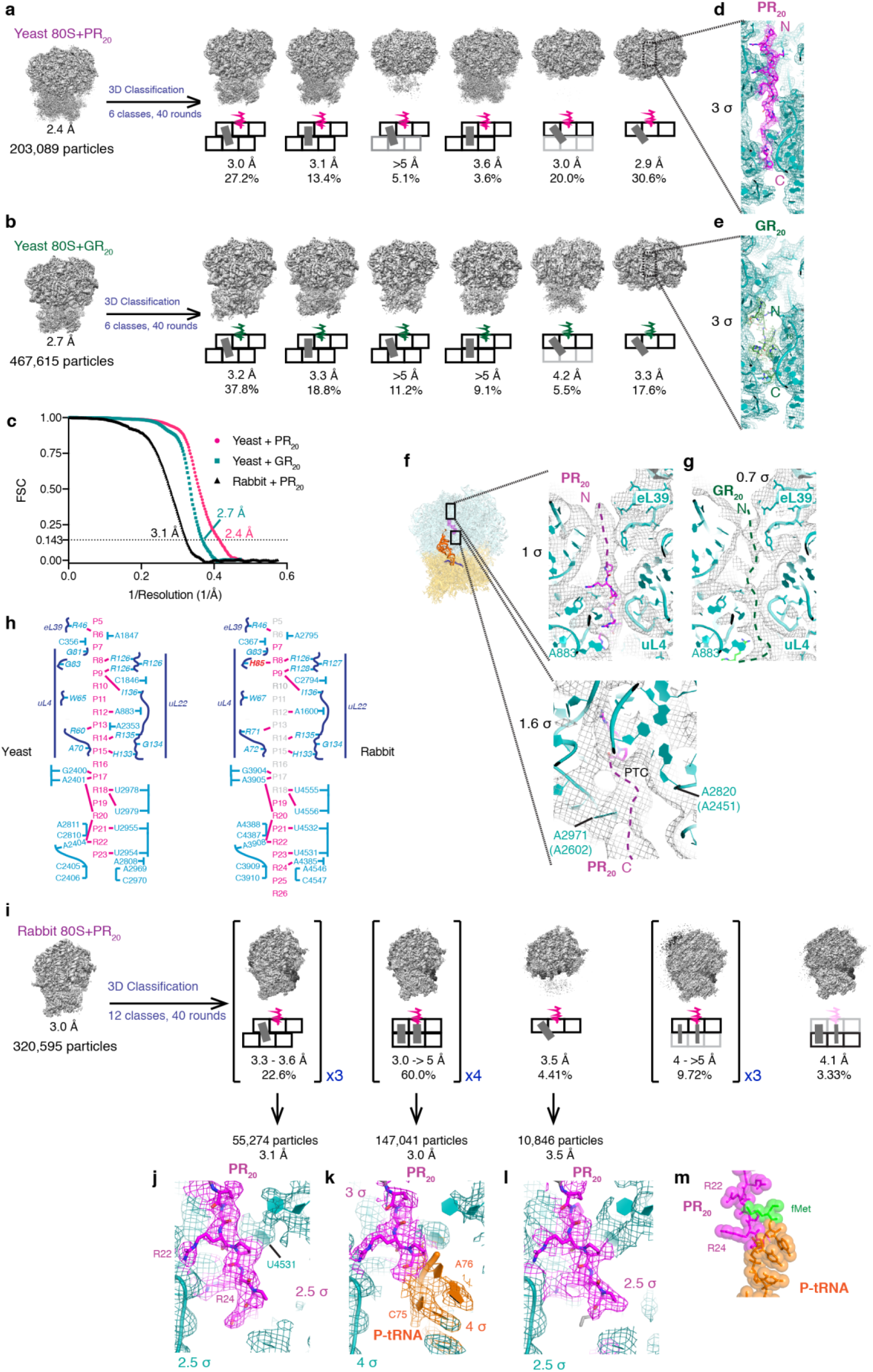
Cryo-EM structures of yeast and rabbit 80S ribosomes with PR_20_ or GR_20_. (**a**) Maximum likelihood classification for the yeast 80S•tRNA^fMet^•PR_20_ dataset shows PR_20_ binding at the polypeptide tunnels of 80S and 60S particles. Cartoons depict the states/conformations of the ribosome as follows: black grid are large (top) and small (bottom) subunits with their respective E (left square), P (middle square), and A (right square) sites. Gray bars are tRNA bound to these sites. Decreased width of tRNA or gray grid indicates partial occupancy. Offset of top and bottom rows indicate rotation of the small subunit relative to the large subunit. Magenta squiggle indicates PR_20_ bound to the polypeptide tunnel. (**b**) Maximum likelihood classification for the yeast 80S•tRNA^fMet^•GR_20_ dataset shows GR_20_ binding at the polypeptide tunnels of 80S and 60S particles. Maps are shown at 3 σ; cartoon schematics are as in (**a**), except that green squiggle indicates GR_20_ bound to the polypeptide tunnel. (**c**) Fourier shell correlation (FSC) curves for cryo-EM maps used for model building and structure refinements. (**d**) Cryo-EM density for PR_20_ bound to the yeast 60S ribosomal subunit. Cryo-EM map is shown at 3 σ. (**e**) Cryo-EM density for GR_20_ bound to the yeast 60S ribosomal subunit. Cryo-EM map is shown at 3 σ. (**f**) The yeast 80S ribosome with PR_20_ contains lower-resolution features extending away from the modeled density towards 60S surface (top panel) and from the peptidyl transferase center (PTC) to intersubunit space (bottom panel). The cryo-EM map was low-pass filtered to 5 Å, and a B-factor of 100 Å^2^ was applied and the mesh is shown at 1 σ and 1.6 σ for the top and bottom panel, respectively. (**g**) The yeast 80S ribosome with GR_20_ contains lower-resolution features extending away from the modeled density towards 60S surface. The cryo-EM map was low-pass filtered to 5 Å, and a B-factor of 100 Å^2^ was applied and is shown at 0.7 σ. (**h**) The ribosomal residues of the polypeptide tunnel (blue) that interact with PR_20_ (magenta) are conserved between yeast (left) and rabbit (right) with one exception (red, a glycine residue in yeast uL4 is a histidine in rabbit uL4). Protein residues are shown in italics, rRNA are regular print. Nucleobase and side chain interactions are shown by pink dashes, backbone interactions are shown by proximity. (**i**) Maximum likelihood classification for the rabbit 80S•tRNA^fMet^•PR_20_ dataset reveals 80S maps with different small-subunit rotation and tRNA occupancy, and 60S maps, yet all classes contain PR_20_ (magenta) in the polypeptide tunnel. (**j**) Density for PR_20_ in the cryo-EM map for the hybrid-state (rotated) 80S particles with P/E tRNA (3 σ). (**k**) Densities for PR_20_ (3 σ) and P-site tRNA (4 σ) are in close contact in the cryo-EM map of the classical (non-rotated) rabbit 80S ribosome bound with P-site tRNA. PR_20_ is displaced down the tunnel, compared to that in the absence of P-site tRNA. An aminoacyl (see **m**) or peptidyl moiety at A76 of the P-site tRNA would sterically clash with PR_20_. (**l)** Density for PR_20_ in the cryo-EM map of rabbit 60S particles is similar to that in the 80S particles (see **j**). (**m**) Amino-acylated P-site tRNA (fMet shown in green spheres, from PDB:5YUM) would sterically clash with PR_20_ in the non-rotated rabbit 80S ribosome.

**Supplementary Fig. 3.**
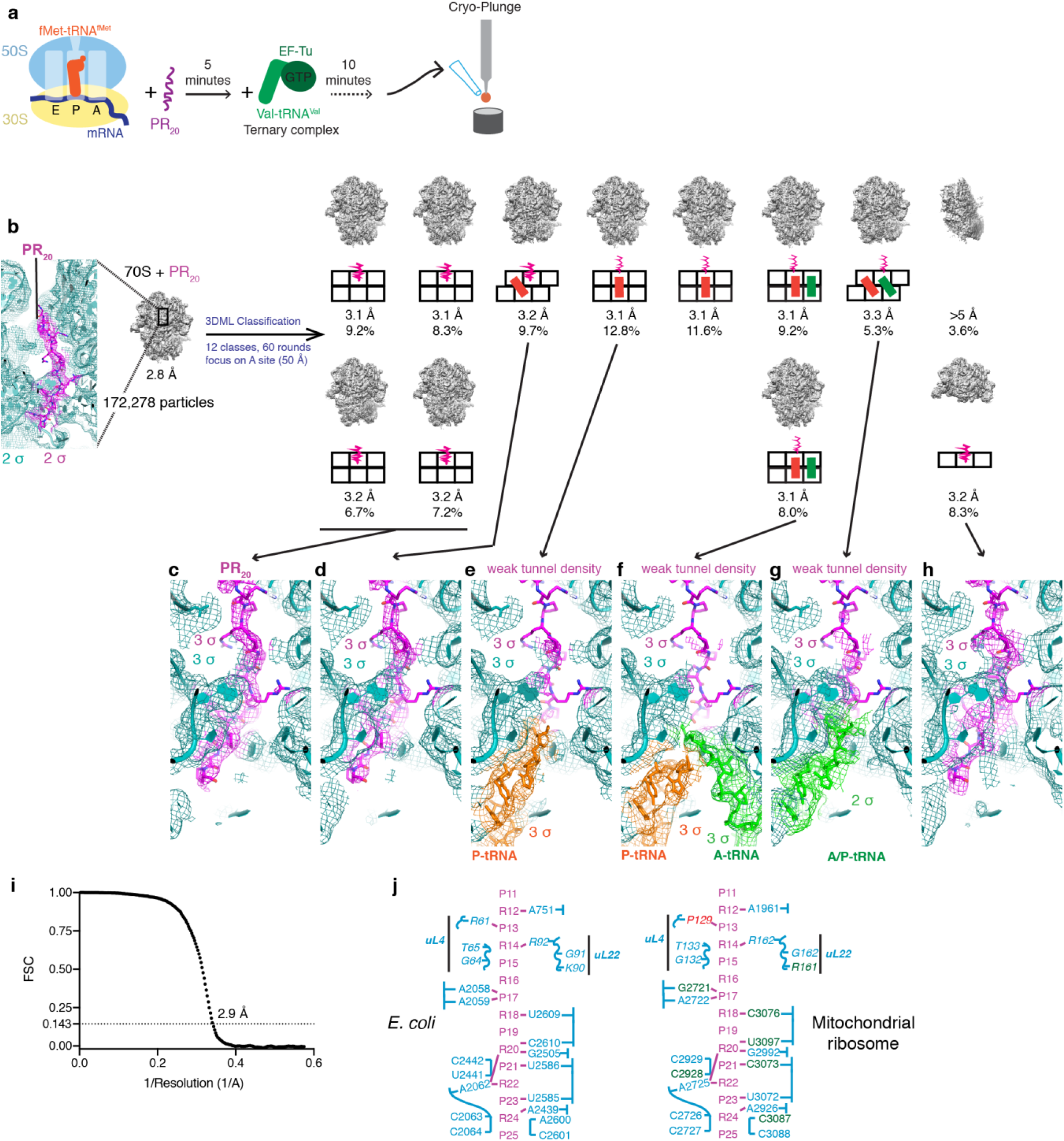
Cryo-EM analyses of *E. coli* 70S ribosomes during elongation in the presence of PR_20_. (**a**) Scheme for a translation elongation reaction on the 70S ribosome with PR_20_, captured by cryo-EM. (**b**) Maximum likelihood classification of the 70S-PR_20_ dataset schematics shown as in **Supplementary Fig. 2a.** (**c)** PR_20_ occupies the polypeptide tunnel of the 70S ribosome. PR_20_ was built into the 70S map containing no tRNA (2.9 Å resolution). The map was B-factor sharpened (−32 Å^2^) and is shown at 3 σ. (**d**) PR_20_ occupies the polypeptide tunnel of the 70S ribosome in the hybrid state with P/E tRNA. Model of the 50S and PR_20_ from (**c**) were rigid-body fit into map of the 70S ribosome with rotated 30S subunit bound with P/E tRNA. Map is shown, without sharpening, at 3 σ. (**e**) Density in the tunnel is weak in the presence of fMet-tRNA (orange). 50S and PR_20_ from (**c**) were rigid-body fit into the map of 70S with P- and E-site tRNAs. P-site tRNA (fMet-tRNA^fMet^) from PDB: 5UYM (Loveland et al., 2017) is shown. (**f**) Density in the tunnel is weak in the presence of both the A-site (green) and P-site (orange) tRNAs. Deacylated P-tRNA (P site) and dipeptidyl-tRNA (A site) from PDB: IVY5 (Polikanov et al., 2014) are shown. (**g**) Density in the tunnel is weak in the presence of the product dipeptidyl-tRNA (A/P hybrid state; green). Analog of dipeptidyl-A/P-tRNA from PDB:1M90 (Hansen et al., 2002) is shown. (**h**) Strong density for PR_20_ in the polypeptide tunnel of the 50S ribosomal subunit. (**i**) Fourier shell correlation (FSC) curve for the 70S map used to model the 70S•PR_20_ structure (see panel **c**). (**j**) Comparison of the *E. coli* polypeptide tunnel with that of the human mitochondrial ribosome. PR residues from the *E. coli* cryo-EM structure (this work) are shown (pink). Interacting ribosomal protein residues are shown in italics, rRNA are regular print (blue): nucleobase and side chain interactions are shown by pink dashes, backbone interactions are shown by proximity. Conservative differences (ex. purine/purine) are green, non-conservative difference R61/P129 is red.

**Supplementary Fig. 4.**
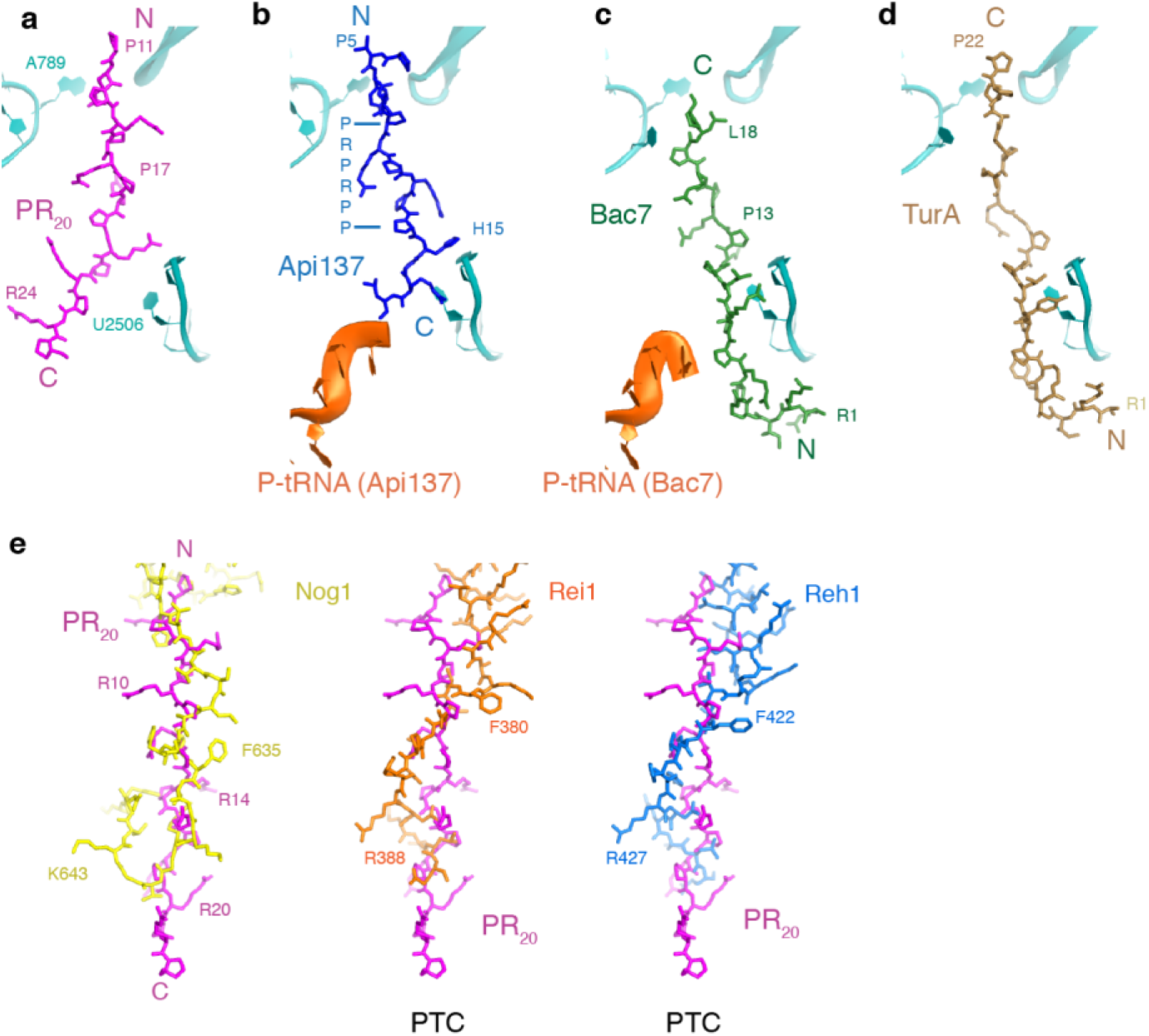
Binding sites of PR_20_ and GR_20_ overlap with those of eukaryotic proline-rich antimicrobial peptides (PrAMPs) and 60S ribosomal subunit biogenesis factors. (**a-d**) Comparison of the position of PR_20_ in the 70S tunnel (**a**, magenta; this work) with those of eukaryotic proline-rich antimicrobial peptides (PrAMPs) bound to bacterial 70S ribosomes: (**b**) Api137 (blue; PDB:5O2R (Florin et al., 2017b)) (**c**) Bac7 (green; PDB:5HAU (Gagnon et al., 2016a)), (**d**) TurA (brown; PDB: 6FKR (Mardirossian et al., 2018). Ribosomal protein uL22 and helices 35a and 90 of 23S rRNA of the polypeptide tunnel are shown in cyan for reference. (**e**) Cryo-EM structures of yeast 60S biogenesis intermediates bound with Nog1 (yellow), Rei1 (orange) and Reh1 (blue) (PDB: 3JCT (Wu et al., 2016), PDB:5APN (Greber et al., 2016) and PDB:5H4P (Ma et al., 2017)) show that the binding sites overlap with that of PR_20_ in the yeast 60S subunit (magenta; this work).

## Supplementary Information

### Cryo-EM classification suggests how PR_20_ interferes with peptidyl transfer on E. coli 70S ribosomes

To understand how PR_20_ structurally interferes with dipeptide bond formation, we have reconstituted a 70S elongation ribosome with authentic aminoacyl-tRNA substrate in the P site and cognate ternary complex EF-Tu•GTP•Val-tRNA^Val^ to deliver aminoacyl-tRNA to the A site for peptidyl transfer (**Supplementary Fig. 3a**). The Cryo-EM data set was collected and analyzed using 3D maximum likelihood classification in cisTEM (**Fig. 3b-c**). The classification separated the empty ribosomes (no tRNA), from the substrate (vacant A site) and product (tRNA-dipeptide-bound A-site) states (**Supplementary Fig. 3b**). In the presence of PR_20_, the non-functional empty 70S ribosome were abundant suggesting a defect in initiation and peptide bond formation.

The occupancy of the polypeptide tunnel by PR_20_ correlates with the occupancy of the peptidyl transferase center by tRNAs (**Supplementary Fig. 3c-h**), consistent with direct inhibition of peptidyl transfer. The DPR density is best resolved in the polypeptide tunnel of the ribosomes without tRNA (**Fig. 3b, Supplementary Fig. 3c-d**). Similar to the 80S•tRNA•PR_20_ complex, lower resolution regions flanked well-ordered regions of poly-PR density at the polypeptide tunnel constriction, coinciding with the macrolide antibiotics’ binding site (**Fig. 3c**). In the presence of tRNA, however, the density in the tunnel and tunnel constriction is very weak (**Supplementary Fig. 3e-g**), suggesting partial occupancy or a shift of poly-PR away from the P-site cleft and into the tunnel. Poly-PR displacement in reaction intermediates suggests that the DPR protein may sample multiple registers along the polypeptide tunnel slowing extension of nascent polypeptides towards the constriction. Nevertheless, strong density at the peptidyl transferase center suggests that the main mode of inhibition is via direct competition with tRNA binding to both the P site (during initiation and elongation) and the A site (during initial round(s) of elongation).

---

**Table S1.** Cryo-EM data collection, refinement and validation statistics

|  | S.c. 80S<br>poly-GR | S.c. 80S<br>poly-PR | O.c. 80S<br>poly-PR | E. c. 70S<br>poly-PR |
| --- | --- | --- | --- | --- |
| **Data collection and processing** |  |  |  |  |
| Magnification | 47,619x | 47,619x | 57,471x | 57,471x |
| Voltage (kV) | 300 | 300 | 200 | 200 |
| Electron exposure (e<sup>-</sup>/Å<sup>2</sup>) | 30 | 30 | 30 | 35 |
| Defocus range (μm) | 0.4-5 | 0.4-5 | 0.4-3 | 0.4-5 |
| Pixel size (Å) | 1.05 | 1.05 | 0.87 | 0.87 |
| Symmetry imposed | C1 | C1 | C1 | C1 |
| Initial particle images (no.) | 467,615 | 203,089 | 320,595 | 172,278 |
| Final particle images (no.) | 467,615 | 203,089 | 63,475 | 49,219 |
| Map resolution (Å)** | 2.7 | 2.4 | 3.1 | 2.9 |
| FSC threshold | 0.143 | 0.143 | 0.143 | 0.143 |
| Map resolution range (Å) | 2.5 - >8 | 2.2 - >8 | 2.8 - >8 | 2.5 - >8 |
| **Refinement** |  |  |  |  |
| Initial model used (PDB code) | 4V88 | 4V88 | 6R5Q | 5UYM |
| Model resolution (Å)* | 2.7 | 2.4 | 3.1 | 2.9 |
| FSC threshold | 0.143 | 0.143 | 0.143 | 0.143 |
| Model resolution range (Å) | 2.5 - >8 | 2.2 - >8 | 2.8 - >8 | 2.5 - >8 |
| Correlation Coefficient (cc_mask)* | 0.89 | 0.87 | 0.87 | 0.84 |
| Real-space R-factor † | 0.19 | 0.19 | 0.18 | 0.20 |
| Map-sharpening B factor (Å<sup>2</sup>) | -25 | -32 | -22 | -30 |
| Model composition* |  |  |  |  |
| Non-hydrogen atoms | 123,855 | 123,892 | 136,560 | 145,570 |
| Protein residues | 6,220 | 6,222 | 6,798 | 6,104 |
| RNA residues | 3,484 | 3,484 | 3,806 | 4,562 |
| B factors (Å<sup>2</sup>)* |  |  |  |  |
| Protein | 69.6 | 71.2 | 131.1 | 133.5 |
| RNA | 77.6 | 66.8 | 126.0 | 114.6 |
| R.m.s. deviations*§ |  |  |  |  |
| Bond lengths (Å) | 0.007 | 0.007 | 0.006 | 0.009 |
| Bond angles (°) | 1.0 | 1.0 | 0.9 | 1.0 |
| Validation |  |  |  |  |
| MolProbity score | 1.88 | 1.93 | 2.16 | 2.25 |
| Clashscore | 6.94 | 6.80 | 11.0 | 15.1 |
| Poor rotamers (%) | 0.7 | 1.2 | 0.6 | 0.8 |
| Ramachandran plot |  |  |  |  |
| Favored (%) | 91.7 | 92.0 | 87.8 | 89.2 |
| Allowed (%) | 8.0 | 7.7 | 10.7 | 9.7 |
| Disallowed (%) | 0.3 | 0.3 | 1.5 | 1.1 |
| Validation (RNA) |  |  |  |  |
| Good sugar pucker (%) | 98.8 | 98.9 | 98.7 | 99.4 |
| Good backbone (%)# | 87.1 | 87.2 | 84.8 | 85.2 |

\*\* from FREALIGN (FSC\_part)
\* from PHENIX
† from RSREF
§ root mean square deviations

